# Hif-1alpha induced expression of Il-1beta protects against mycobacterial infection in zebrafish

**DOI:** 10.1101/306506

**Authors:** Nikolay V. Ogryzko, Amy Lewis, Heather L. Wilson, Annemarie H. Meijer, Stephen A. Renshaw, Philip M. Elks

## Abstract

Drug resistant mycobacteria are a rising problem worldwide. There is an urgent need to understand the immune response to TB to identify host targets that, if targeted therapeutically, could be used to tackle these currently untreatable infections. Here, we use an Il-1β fluorescent transgenic line to show that there is an early innate immune pro-inflammatory response to well-established zebrafish models of inflammation and *Mycobacterium marinum* (Mm) infection. We demonstrate that host-derived hypoxia signalling, mediated by the Hif-1α transcription factor, can prime macrophages with increased levels of Il-1β in the absence of infection, upregulating neutrophil antimicrobial nitric oxide production, leading to greater protection against infection. Our data link Hif-1α to proinflammatory macrophage Il-1β transcription *in vivo* during early mycobacterial infection and importantly highlight a host protective mechanism, via antimicrobial nitric oxide, that decreases disease outcomes and that could be targeted therapeutically to stimulate the innate immune response to better deal with infections.

## Introduction

Pulmonary tuberculosis (TB) is a major world health problem caused by the bacillus *Mycobacterium tuberculosis (Mtb)* (World Health Organization, 2016). It is a current priority for infectious disease research due to increasing rates of multi- and totally-drug resistant strains causing high levels of mortality, especially in the immunocompromised (Koul *et al*, 2011). Mycobacteria are specialised at evading killing mechanisms of the immune system to survive. Mycobacteria and immune cells create a highly organised niche, called the granuloma, in which Mtb can proliferate or enter a latent phase, protected from the immune system (Podinovskaia *et al*, 2013; Ramakrishnan, 2012). In human Mtb infection, bacteria first encounter cells of the innate immune system in and around the lungs, either macrophages in the alveolar space or neutrophils in the surrounding capillary vasculature, before the involvement of adaptive immunity and granuloma formation (Lerner *et al*, 2015; Jasenosky *et al*, 2015). These initial phagocytosis events are followed by the attraction of other innate immune cells which signal to draining lymph nodes to activate the adaptive immune response, signs of which only become apparent 3 to 8 weeks after infection in humans (Jasenosky *et al*, 2015). Although granuloma formation is reasonably well characterised, the initial interactions of the bacteria with the host innate immune cells are less well defined *in vivo.*

Mtb, like many other bacterial and pathogenic microbes, triggers a pro-inflammatory immune response via the activation of TLRs (Toll-like receptors) (Mortaz *et al*, 2015). The activation of the innate immune cells via TLR signalling is a critical early host response to many invading pathogens for successful clearance of infection and, in the absence of TLR signalling, mycobacteria grow unchecked to cause systemic infection (van der Vaart *et al*, 2013). Although mycobacteria can hijack host leukocytes to create a niche for their growth, in zebrafish models many of the initial Mm inoculum are neutralised by macrophages and neutrophils before infection can take hold (Hosseini *et al*, 2016; Cambier *et al*, 2013). Early mycobacterial interaction with host leukocytes is critical for the pathogen, and manipulation of the macrophage by the bacteria is required for establishment of a permissive niche in which the bacteria can grow and build its host derived protective structure, the granuloma (Meijer, 2016; Guirado *et al*, 2013). Indeed the control of the macrophage by Mm may happen early in infection, as there is a phase of infection from 6 hours to 1 day post infection in the zebrafish model that is characterised by a dampening of the cytokine transcriptional response (Benard *et al*, 2016). Greater understanding of the diverse phenotype of macrophages immediately after infection may allow therapeutic tuning to provide maximal early control of mycobacteria during infection (McClean & Tobin, 2016; Dorhoi & Kaufmann, 2015). Recent studies in optically translucent zebrafish infection models have indicated that initial interactions between Mm and macrophages and neutrophils are more complex than originally thought, with successive rounds of bacterial internalisation and leukocyte cell death leading to granuloma formation (Hosseini *et al*, 2016; Cambier *et al*, 2017; Elks *et al*, 2015). The immune molecular mechanisms involved in these early processes are poorly understood.

We have previously demonstrated in a zebrafish/Mm model of TB that the initial immune response to infection can be enhanced by stabilising host-derived Hif-1α (hypoxia inducible factor-1 alpha), leading to reduced bacterial burden (Elks *et al*, 2013). Hif-1α is a major transcriptional regulator of the cellular response to hypoxia, that has been implicated in the activation of macrophages and neutrophils during infection and inflammatory processes (Cramer *et al*, 2003; Elks *et al*, 2011). Stabilisation of Hif-1α in zebrafish upregulated pro-inflammatory neutrophil nitric oxide (NO) production leading to lower mycobacterial burden (Elks *et al*, 2013, 2014a). The mechanisms by which pro-inflammatory cytokines associated with this NO increase are regulated by Hif-1α signalling is not known.

IL-1β is a critical macrophage-derived activator of immune cells with wide-ranging and complex effects on immune signalling and downstream pathways. IL-1β has been shown to be upregulated in the onset and formation of Mm and Mtb granulomas (Di Paolo *et al*, 2015; Bourigault *et al*, 2013; Novikov *et al*, 2011). We hypothesised that IL-1β would be activated in specific immune cell populations early in Mm infection, (within 1-day post-infection, pre-granuloma formation) and that Hif-1α acts via altered expression of this important pro-inflammatory mediator to confer protection against mycobacterial infection. Here, using the zebrafish Mm model and fluorescent transgenic lines, we show that that *il-1β* is transcriptionally upregulated in macrophages early during *in vivo* infection. Stabilisation of Hif-1α upregulates *il-1β* transcription in macrophages in the absence of infection. *il-1β* signalling is required for protective NO production by neutrophils and a subsequent decrease in infection. Our data indicate that protective Hif-1α mediated NO is at least partially regulated by the key pro-inflammatory mediator Il-1β, increasing our understanding of the mechanism of action of the potential therapeutic target, Hif-1α, as a host-derived factor in TB.

## Results

### *il-1β:GFP* is upregulated in macrophages during early and later stage Mm infection

Il-1β is a major macrophage-derived pro-inflammatory cytokine that is upregulated in both inflammation and infection. The initial phase of Mm infection in zebrafish is characterised by a short period of greatly increased pro-inflammatory signalling (before 1 day post-infection, dpi) where the immune system reacts to Mm infection. This is followed by a lag-phase of decreased activity which allows for granuloma formation between 2-3dpi, before cytokine levels rise again in formed larval granulomas at 4dpi (Benard *et al*, 2016; Hosseini *et al*, 2016). However, the levels of pro-inflammatory cytokines have only been previously studied at a transcriptional level in whole embryos or FACS sorted cells, rather than detecting levels *in situ*, over time, in an intact organism (Benard *et al*, 2016).

We hypothesised that Il-1β is a major pro-inflammatory cytokine that would be upregulated by both mycobacterial infection and Hif-1α stabilisation. We have previously shown upregulation of *il-1β* message after induction of inflammation via tailfin transection by qPCR and wholemount *in situ* hybridisation (WISH) in the zebrafish (Ogryzko *et al*, 2014a). *il-1β* is one of the most readily detectable pro-inflammatory cytokines during early granuloma stages of Mm infection and at 1 dpi (Figure 1A) (Van Der Vaart *et al*, 2014). At 1dpi transcription is upregulated 1.7 fold measured by qPCR, compared to PVP injection controls (Figure 1A). Macrophage expression of *il-1β* is greatly under-represented measured in this way on the wholebody level due to the small proportion of cells that contribute to the immune lineage. Therefore, to investigate *il-1β* expression on a cellular level *in vivo*, we developed a BAC (bacterial artificial chromosome) derived *il-1β* promoter driven GFP line, *TgBAC(il-1 β:GFP)SH445*, to assess *il-1β* expression in real-time during mycobacterial infection. We sought to examine *il-1β:GFP* expression in our well-established inflammation assay before investigating its expression during mycobacterial infection. Both wholemount *in situ* hybridisation (WISH) of *il-1β* and *il-1β:GFP* do not exhibit any immune cell expression under basal conditions (Figure S1A-B and Figure 1B). *il-1β:GFP* recapitulates *il-1β* WISH expression in response to tail transection, with upregulation observed in cells in and around the caudal haematopoietic region (CHT), consistent with immune cell expression, (Figure S1A and B) (Ogryzko *et al*, 2014a), although, as expected, the synthesis of GFP occurs over a longer timescale than that of *il-1β* mRNA detected by WISH. Neutrophils are the first cells to respond to tailfin transection with increased *il-1β:GFP*, with fluorescence first observed at 1hpw (hours post-wounding) and still present at 6hpw (Figure S1C). Having demonstrated that the il-1β:GFP is responsive to inflammation in similar cells over a similar timespan as the *in situ* hybridisation, we sought to investigate its regulation during mycobacterial infection.

**Figure 1.**
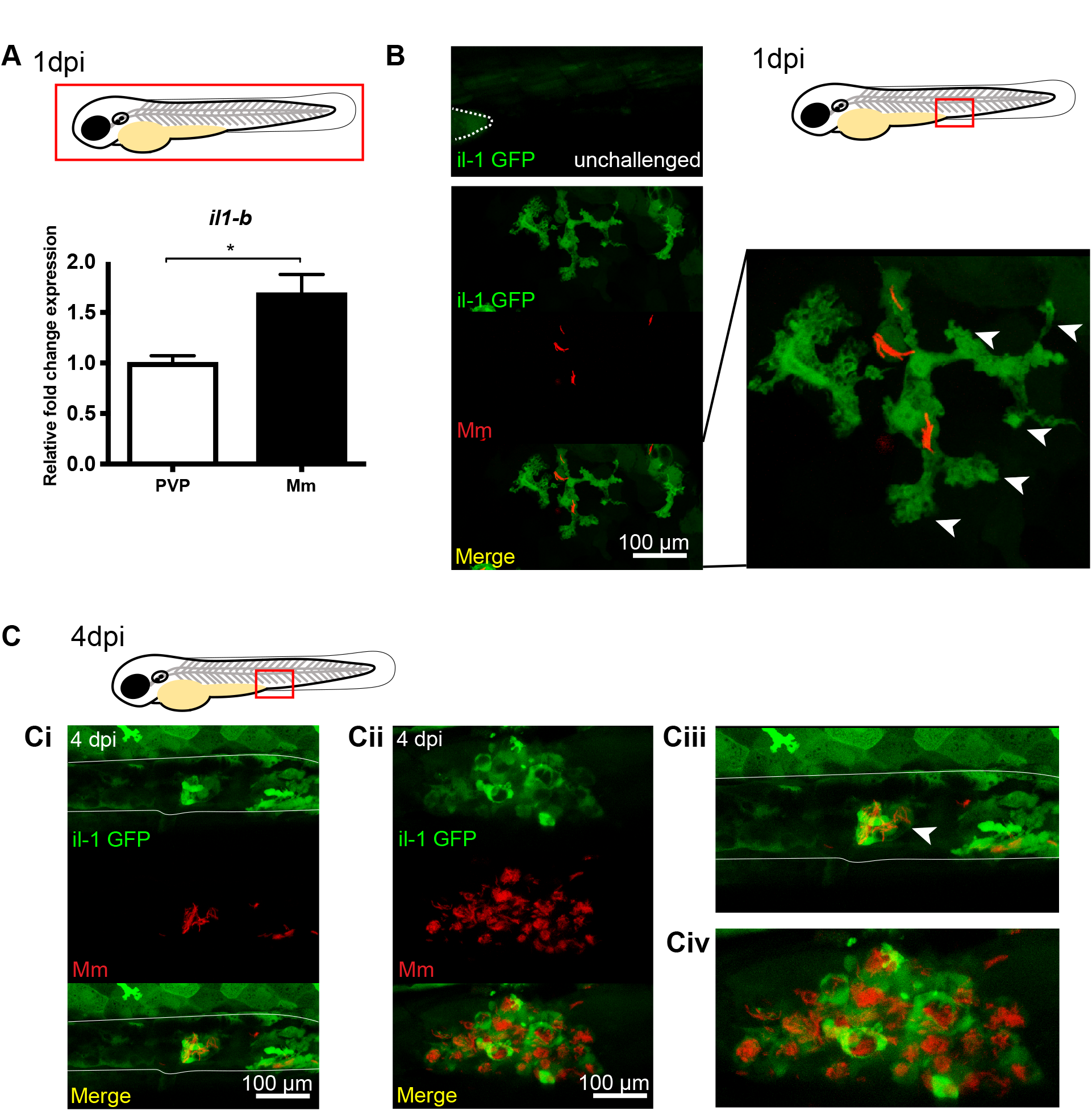
*TgBAC(il-1β:GFP)sh445* is upregulated by Mm in infected macrophages at early and later stage infection. A) Graph showing relative wholebody *il-1β* mRNA expression by SYBRgreen qPCR in Mm infected 1dpi larvae (Mm) and mock injected controls (PVP). Data shown are mean ± SEM, n=3 independent experiments. (B) Fluorescent confocal micrographs of 1dpi larvae, prior to granuloma formation. Unchallenged *TgBAC(il-1 β:GFP)sh445* has no detectable expression in immune cells and low detectable levels in the yolk (dotted line) and some muscle cells. *il-1β* expression was detected by GFP levels, in green, using the *TgBAC(il-1:eGFP)sh445* transgenic line. Mm mCherry is shown in the red channel. Increased levels of *il-1β:GFP* expression were detectable in cells associated with infection. Infected macrophages with *il-1β-GFP* levels have an activated, branched phenotype (white arrowheads). (C) Fluorescent confocal micrographs of 4dpi larvae. *il-1β* expression was detected by GFP levels, in green, using the *TgBAC(il-1:eGFP)sh445* transgenic line. Mm mCherry is shown in the red channel. Increased levels of *il-1β:GFP* expression were detectable in immune cells that are in the blood vessels (Ci and blown up in Ciii, blood vessel indicated by solid white lines) and in early tissue granulomas (Cii and blown up in Civ).

We used the *TgBAC(il-1 β:GFP)sh445* line to show that GFP is expressed in cells proximal to Mm infection sites at pre-granuloma phases (1dpi) (Figure 1B) and in larval granulomas (4dpi) (Figure 1C). Many of these cells contained Mm and had the appearance of activated immune cells with a dynamic branched phenotype (Figure 1B and Movie S1). The earliest timepoint at which *il-1β:GFP* could be detected by confocal microscopy was between 6 and 8 hours post infection, (Figure 2A), consistent with rapid transcriptional activation of the *il-1β* promoter after infection and similar to the timing of macrophage *il-1β:GFP* expression after tailfin transection (Figure 2B). *il-1β:GFP* was predominantly upregulated in infected macrophages at 1dpi (Figure 2C) consistent with their containment of phagocytosed Mm (Figure 1B). These data demonstrate that, during early stages of infection, *il-1β* is transcriptionally activated in infected macrophages as part of an early pro-inflammatory response.

**Figure 2.**
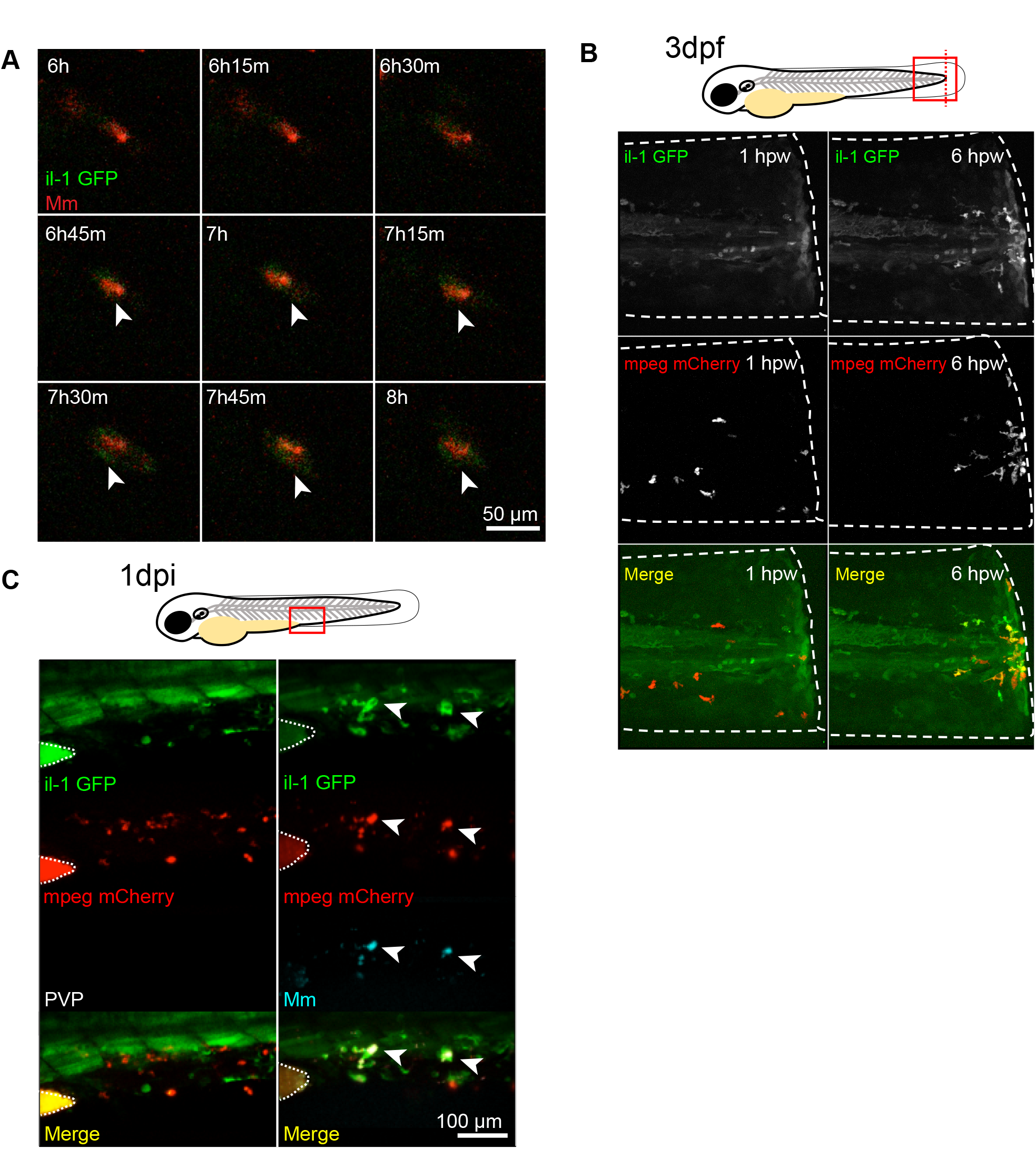
*il-1β:GFP* is activated 6-8 hours after challenge in macrophages. (A) Fluorescent confocal micrographs of a timelapse between 6 to 8 hours post Mm infection. Mm mCherry is shown in the red channel and *il-1β:GFP* in the green channel with the microscope settings set to detect low GFP levels. Arrowheads indicate the emergence of *il-1β:GFP* expression in an infected cell. (B) Fluorescent confocal micrographs of *TgBAC(il-1 β:GFP)sh445* crossed to *Tg(mpeg1:mCherryCAAX)sh378* line labelling macrophages. The tailfin was transected at 3dpf and fluorescence imaging was performed at the wound at 1 hour post wound (1hpw) and 6hpw. Red macrophages are not positive for *il-1β:GFP* expression at 1hpw and the first detectable *il-1β:GFP* expression is found in the macrophages at 6hpw. (C) Fluorescent confocal micrographs of 1dpi caudal vein region of infection. *il-1β* expression was detected by GFP levels, in green, using the *TgBAC(il-1β:eGFP)sh445* transgenic line. Macrophages are shown in red using a *Tg(mpeg1:mCherryCAAX)sh378* line. Mm Crimson is shown in the blue channel (right panels) with a PVP control (left panels). Without infection there is little overlap of *il-1β:GFP* and *mpeg:mCherry*, while in infected larvae macrophages have higher levels of *il-1β:GFP.* Arrowheads indicate infected macrophages with high levels of *il-1β:GFP.* Dotted lines indicate the yolk extension of the larvae where there is nonspecific fluorescence.

### Stabilisation of Hif-1α upregulates *il-1β:GFP* at early stages of infection

We have previously shown that stabilisation of Hif-1α induces neutrophil pro-inflammatory nitric oxide production (Elks *et al*, 2013, 2014a). We hypothesised that this may be a part of an increased pro-inflammatory profile in innate immune cells, therefore we tested whether Hif-1α is upregulating a pro-inflammatory program in the absence of infection using the *il-1β:GFP* transgenic line. Dominant active Hif-1α significantly increased *il-1β:GFP* expression in the absence of Mm infection at 2dpf, while dominant negative Hif-1α caused no difference in *il-1β:GFP* expression (Figure 3A and B). These data indicate that *il-1β* expression is part of a pro-inflammatory response to increased Hif-1α levels that could aid the host response to Mm challenge.

**Figure 3.**
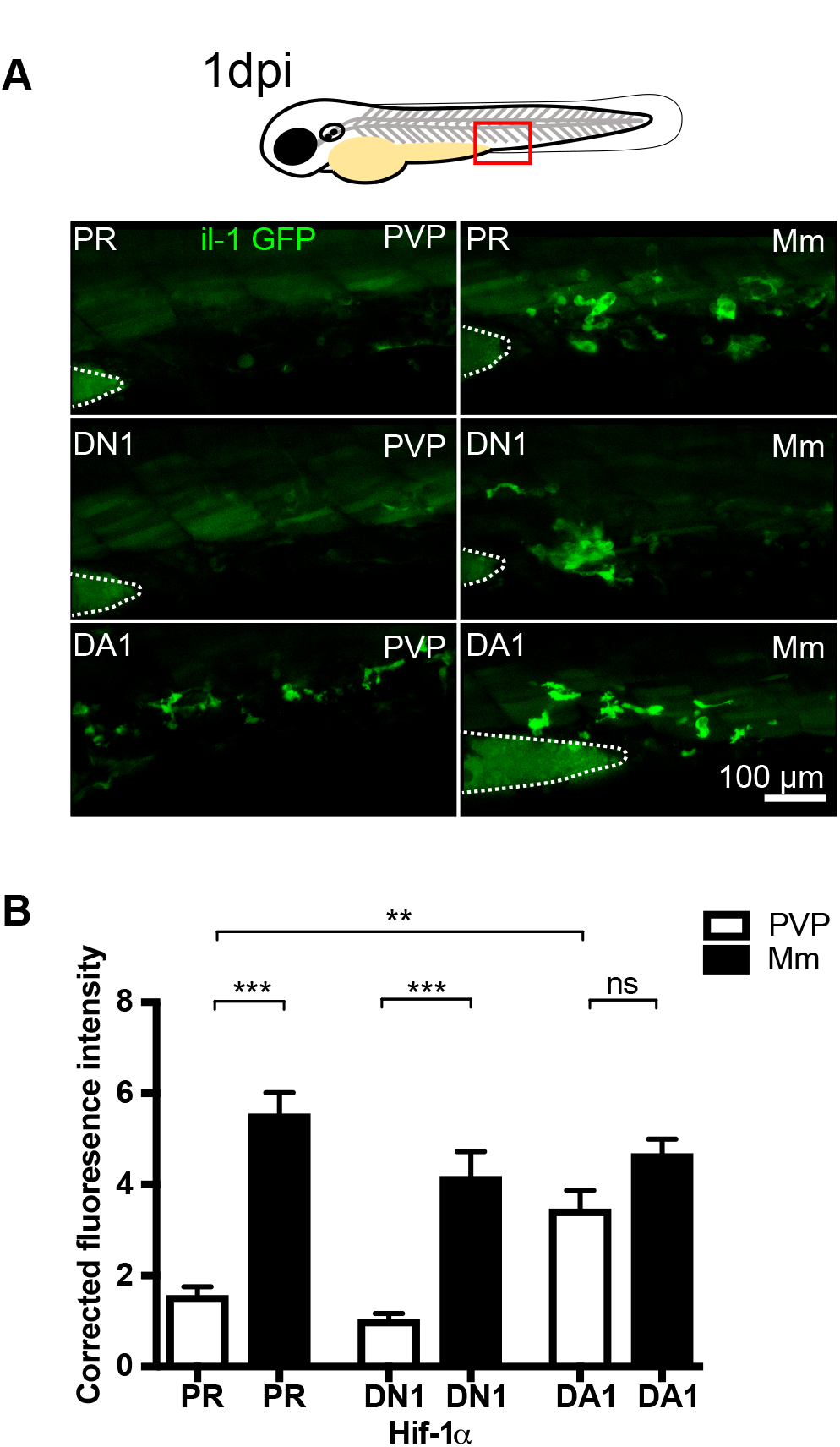
*il-1β:GFP* is upregulated in the absence of infection by DA Hif-1α. (A) Fluorescent confocal micrographs of 1dpi caudal vein region of infection. *il-1β:GFP* expression was detected by GFP levels, in green, using the *TgBAC(il-1:eGFP)sh445* transgenic line. Larvae were injected at the 1 cell stage with dominant negative (DN) or dominant active (DA) Hif-1α or phenol red (PR) control. Noninfected larvae are in the left panels (PVP) and Mm Crimson infected larvae are in the right panels (Mm). Dotted lines indicate the yolk extension of the larvae where there is non-specific fluorescence. (B) Corrected fluorescence intensity levels of *il-1β:GFP* confocal z-stacks in uninfected larvae (PVP, empty bars) and infected larvae (Mm, filled bars) at 1dpi. Dominant active Hif-1α (DA1) had significantly increased *il-1β:GFP* levels in the absence of Mm bacterial challenge compared to phenol red (PR) and dominant negative Hif-1α (DN1) injected controls. Data shown are mean ± SEM, n=24-48 cells from 4-8 embryos representative of 3 independent experiments.

### Inhibition of *il-1β* increases Mm burden and inhibits the Hif-1α nitric oxide response

Il-1β is a major pro-inflammatory cytokine that in many infections is instrumental in coordinating the immune response (Cohen, 2014; Ogryzko *et al*, 2014b). We sought to test whether Il-1β was important in early Mm infection. When Il-1β was blocked using a well-characterised and validated *il-1β* morpholino, the morphants showed significantly increased infection compared to control morphants (Figure 4A and B). We have previously shown that stabilisation of Hif-1α induces pro-inflammatory neutrophil nitric oxide production, via inducible nitric oxide synthase (iNOS) (Elks *et al*, 2013, 2014a). DA Hif-1α was not sufficient to reduce Mm infection levels when *il-1β* expression was blocked (Figure 4A and B) suggesting that the *il-1β* response to Mm infection is critical to control infection. These results were supported by generation of an *il-1β* null mutant (*il-1 β^SH446^/ il-1β^SH446^*) (Figure S2), in which DA Hif-1α also did not decrease infection, while in wildtype siblings infection was reduced (Figure 4C and D).

**Figure 4.**
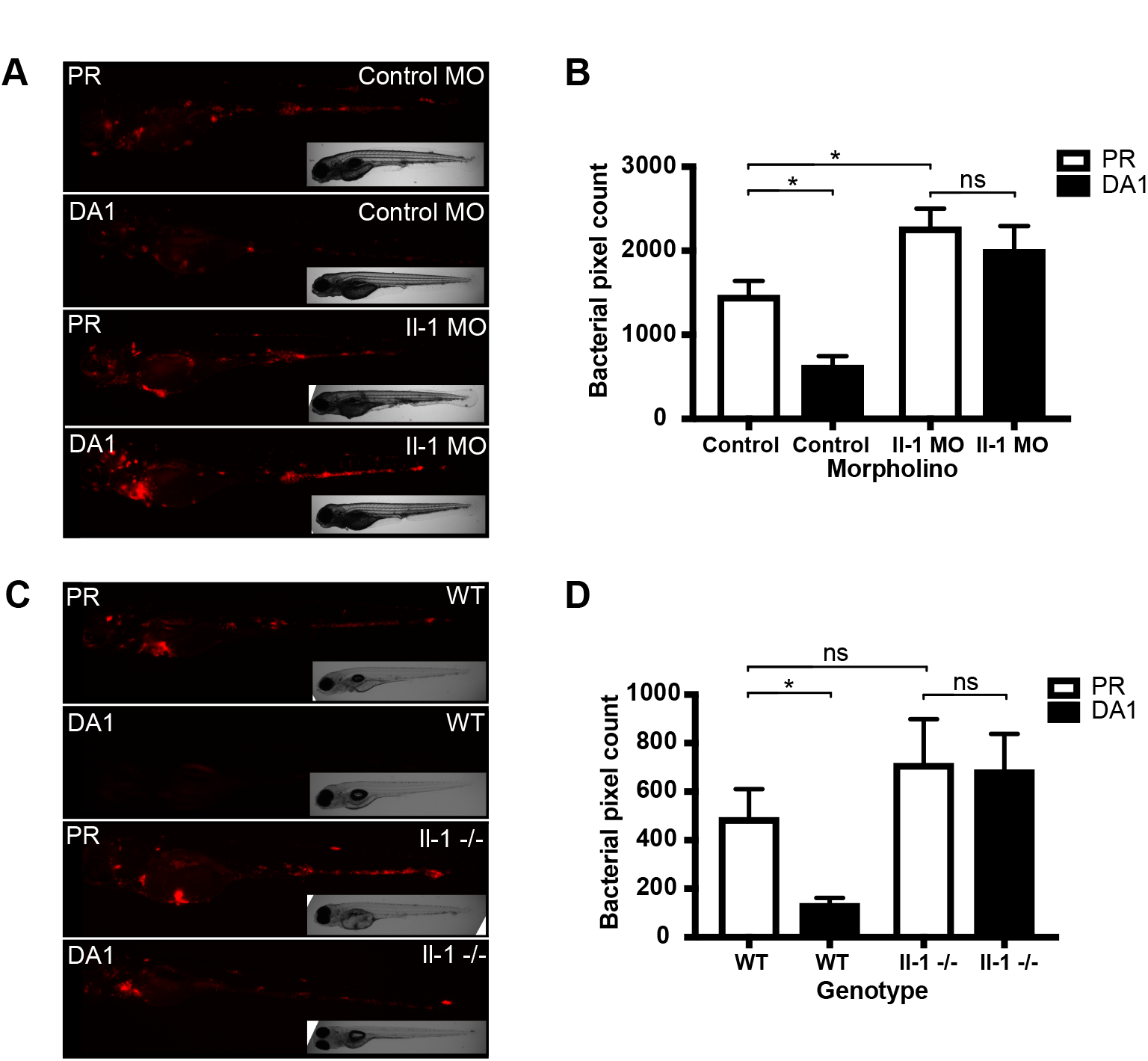
*il-1β* knockdown abrogates the protective effect of DA Hif-1α on bacterial burden. (A) Stereo-fluorescence micrographs of Mm mCherry infected 4dpi larvae after injection with DA Hif-1α (DA1) and the *il-1β* morpholino (Il-1 MO), using the standard control morpholino and phenol red (Control) as a negative control. (B) Bacterial burden of larvae shown in (A). Data shown are mean ± SEM, n=46-50 as accumulated from 3 independent experiments. (C) Stereo-fluorescence micrographs of Mm mCherry infected 4dpi larvae of *il-1β* knockout (il-1-/-) and sibling wildtype controls (WT) after injection of DA Hif-1α (DA1) or phenol red (PR) as a negative control. (D) Bacterial burden of larvae shown in (C). Data shown are mean ± SEM, n=16-20 as accumulated from 3 independent experiments.

NO production is found primarily in neutrophils after Mm infection in zebrafish larvae (Figure S3) (Elks *et al*, 2013, 2014b). We have previously demonstrated that inhibiting production of nitric oxide by Nos2 can block the antimicrobial effect of DA Hif-1α (Elks *et al*, 2013). Blocking Il-1β production also significantly dampened the neutrophil nitric oxide response after Mm infection at 1dpi (Figure 5A and B). As we have previously observed, DA Hif-1α upregulated NO in the absence of infection (PVP) an effect that is dampened by introduction of the bacteria (Mm) through currently unknown mechanisms, (Figure 5C and D) (Elks *et al*, 2013). Here, we find that *il-1β* MO blocked the increased production of nitrotyrosine by DA Hif-1α in the absence of bacteria (PVP) (Figure 5C and D). These results show that Hif-1α activation of Nos2 may, at least in part, be acting through *il-1β* activation (Figure 6) and hint at a much more complex regulation of pro-inflammatory signalling by Hif-1α than simply acting on Hif responsive elements (HREs) in the promoter of Nos2.

**Figure 5.**
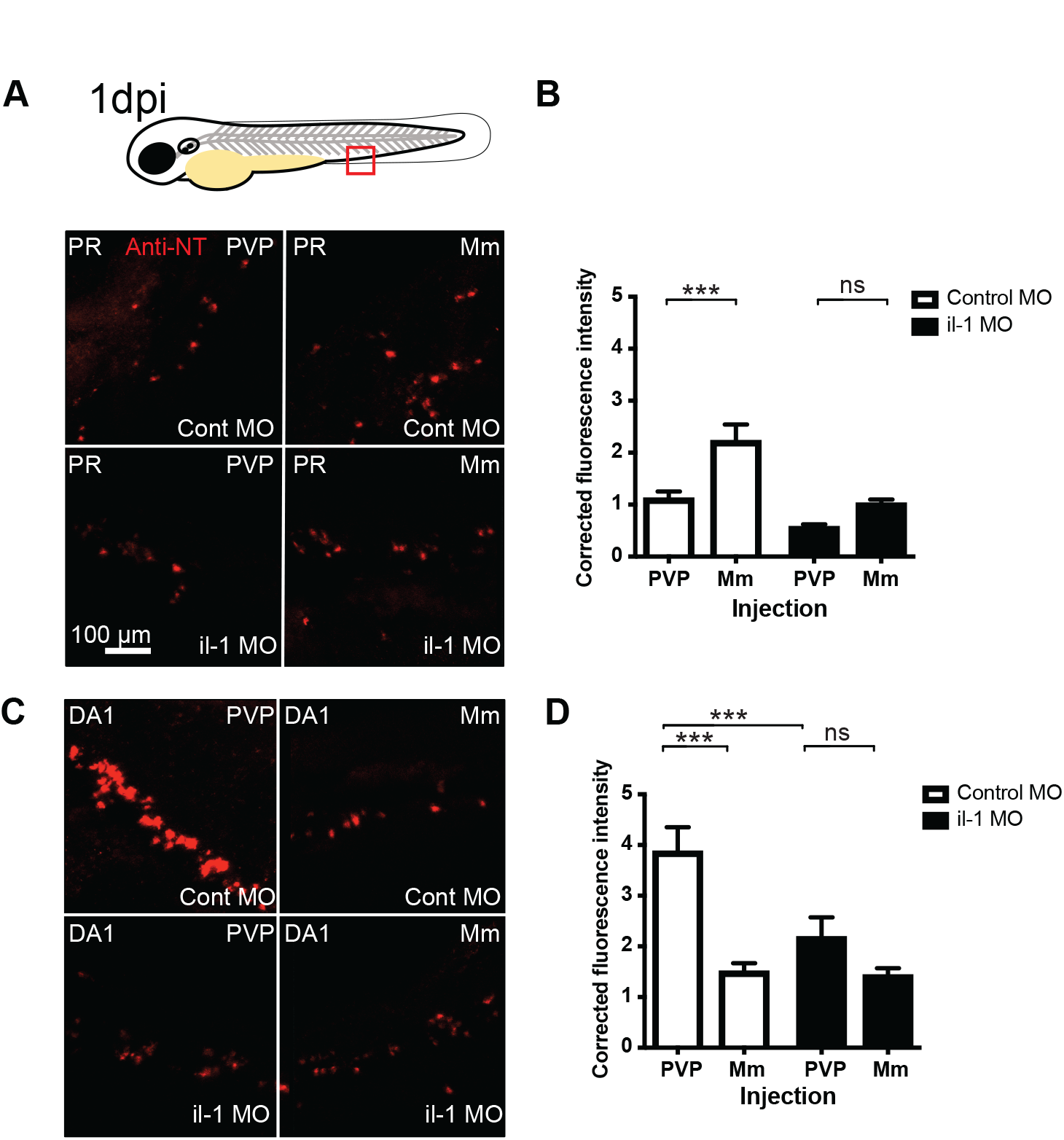
*il-1β* knockdown abrogates DA Hif-1α dependent nitrotyrosine production. (A) Example fluorescence confocal z-stacks of the caudal vein region of embryos stained with Alexa-633 labelled anti-nitrotyrosine antibody (red), imaged at 1dpi in the presence or absence of Mm infection. One-cell stage embryos were injected with phenol red (PR). One-cell stage embryos we co-injected with *il-1β* morpholino or (il-1 MO) or standard control morpholino (Cont MO). At 1dpi larvae were either infected with Mm mCherry (Mm), or PVP as a non-infected control (Mm channel not shown in these panels). (B) Example fluorescence confocal z-stacks of the caudal vein region of embryos stained with Alexa-633 labelled anti-nitrotyrosine antibody (red), imaged at 1dpi in the presence or absence of Mm infection. One-cell stage embryos were injected with dominant active Hif-1α (DA). One-cell stage embryos we co-injected with *il-1β* morpholino or (il-1 MO) or standard control morpholino (Cont MO). At 1dpi larvae were either infected with Mm mCherry (Mm), or PVP as a non-infected control (Mm channel not shown in these panels). (C) Corrected fluorescence intensity levels of anti-nitrotyrosine antibody confocal z-stacks of phenol red (PR) control injected embryos in the presence or absence of Mm infection at 1dpi. Control morpholino is shown in the clear bars and *il-1β* morpholino (il-1 MO) in the filled bars. Data shown are mean ± SEM, n=54-59 cells from 10-12 embryos accumulated from 3 independent experiments. (D) Corrected fluorescence intensity levels of anti-nitrotyrosine antibody confocal z-stacks of dominant active Hif-1α (DA1) injected embryos in the presence or absence of Mm infection at 1dpi. Control morpholino is shown in the clear bars and il-1 morpholino (il-1 MO) in the filled bars. Data shown are mean ± SEM, n=54-59 cells from 10-12 embryos accumulated from 3 independent experiments.

**Figure 6.**
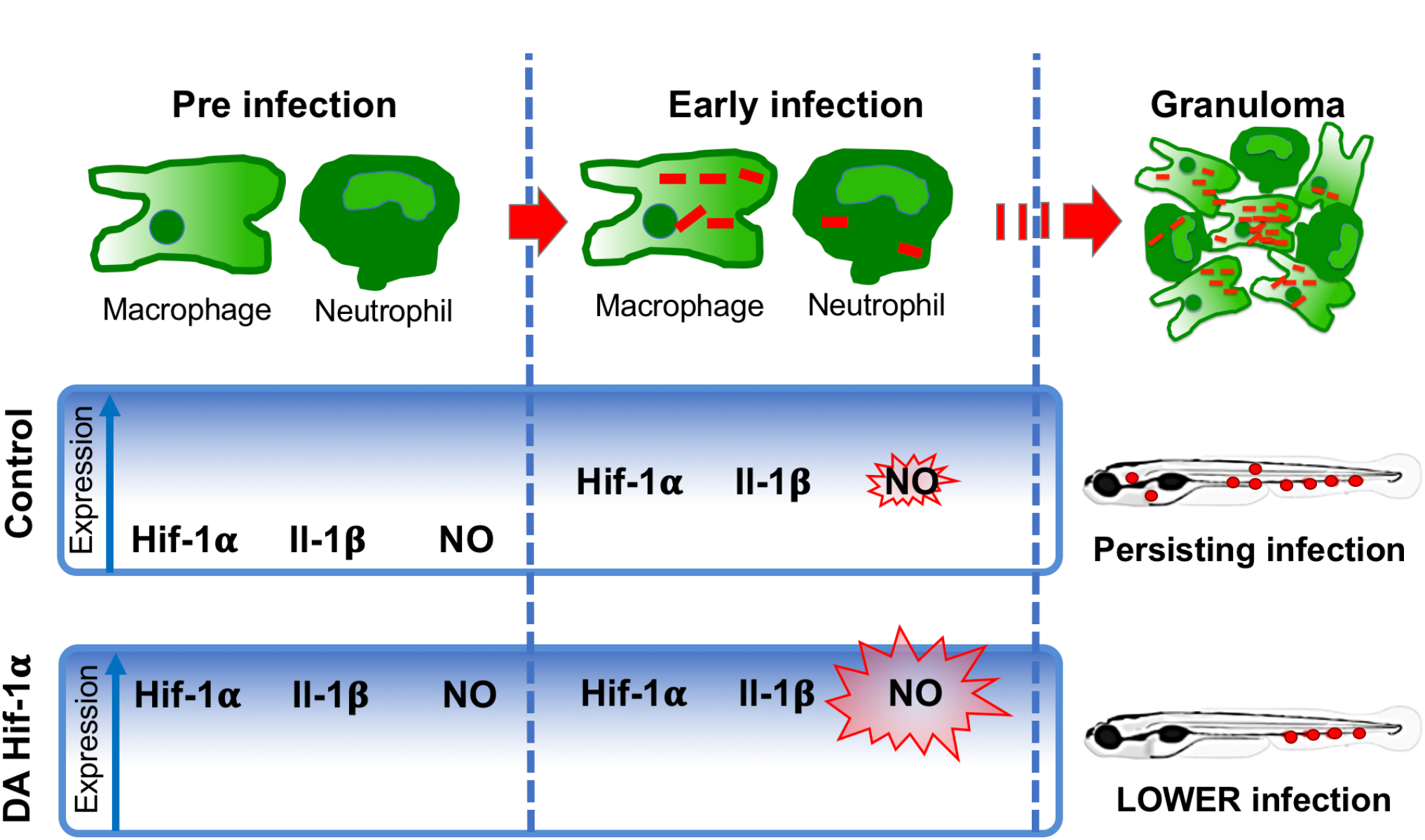
Hif-1α stabilisation leads to upregulation of *il-1β* and increased neutrophil nitric oxide production that is protective against infection. During normal (control) Mm infection Hif-1α, Il-1β and NO transcript levels rise after infection, but are not sufficient to control infection (Elks *et al*, 2013). When Hif-1α is stabilised, Il-1β and subsequent neutrophil NO upregulation occurs in the absence of infection, priming the immune response to better deal with infection leading to lower burden.

## Discussion

Antimicrobial resistance is a rising problem in TB infections worldwide and there is an urgent need to understand the regulation of host-immunity by TB so that we can target host-derived factors to help tackle disease. Our data identify an early pro-inflammatory response, involving macrophage *il-1β* expression, that is important for the onset of early disease, but ultimately fails to control infection leading to granuloma formation. Using a well-established zebrafish Mm model of TB, we show that manipulation of Hif-1α can stimulate this pro-inflammatory network, aiding the host fight against infection, moving towards early clearance of infection. Specifically, we identify that Hif-1α driven Il-1β contributes to the NO response, a response we have previously shown to be host protective (Elks *et al*, 2013, 2014a).

Here, we took advantage of a novel transgenic zebrafish line to understand the dynamics and cell specificity of *il-1β* production in inflammation and mycobacterial infection, with a focus on the understudied early stages (<1dpi) of the innate immune response to TB infection. We confirmed that the *il-1β:GFP* expression of our line was faithful to *il-1β* transcription by following its expression in a well-characterised tailfin transection model of inflammation and comparison to *in situ* hybridisation data (Ogryzko *et al*, 2014a; Renshaw & Loynes, 2006). Furthermore, the expression pattern of our BAC transgenic line closely matches another recently published BAC promoter driven *il-1β* transgenic (Hasegawa *et al*, 2017). The *il-1β:GFP* line also displayed some GFP signal in muscle and epithelial cells in the tail. Similar GFP expression can be seen when driven by NF-kB response elements (Kanther *et al*, 2011) but not by WISH, suggesting this might be off-target expression resulting from the promoter region missing some negative regulatory elements, however, it could also be specific expression that is at too low a level to be detectable by *in situ* hybridisation. Although previous studies have shown *il-1β:GFP* to be upregulated in leukocytes at a tailfin transection (Hasegawa *et al*, 2017), we combined the *il-1β:GFP* line with leukocyte specific transgenics to show that neutrophils are the first to respond at the wound, with macrophages both migrating to and upregulating *il-1β:GFP* at later timepoints.

The Mtb granuloma is widely studied, both in terms of immunohistochemistry of human granulomas, and in mammalian models (Ulrichs & Kaufmann, 2006; Flynn *et al*, 2011; Via *et al*, 2008). These studies have demonstrated that the granuloma is rich in pro-inflammatory cytokine production. This pro-inflammatory environment has been observed in human TB, with IL-1β found to be in high levels in pleural fluid from TB patients with granulomas present (Orphanidou *et al*, 1996). Here we observe that the pro-inflammatory response is present at pre-granuloma stages. Lack of a pro-inflammatory response has been linked to poor treatment outcomes indicating that this host response is important even in the presence of antimycobacterial agents (Waitt *et al*, 2015). The upregulation of proinflammatory cytokines in mycobacterial infection has also been shown in the zebrafish/Mm larval model of TB granulomas, but previous studies have mainly relied on immunohistochemistry and/or transcriptomics data from either wholebody larvae or FACs sorted immune cell populations (Benard *et al*, 2016; Marjoram *et al*, 2015). Using live cell imaging we found that *il-1β* transcription was upregulated at the granuloma formation stage, however we also demonstrated that it is upregulated before the granuloma stage within 6-8 hours hpi. Upon infection *il-1β:GFP* expression was predominantly upregulated in infected macrophages indicating that within the first 24 hours of infection there is a macrophage pro-inflammatory response. Murine and human cell studies have indicated that macrophages are able to produce IL-1β a few hours after mycobacterial challenge indicating that an early response is also present in mammalian systems, at least on a cellular level (Di Paolo *et al*, 2015; Robinson & Nau, 2008). Our observations are in line with our previous observation of Hif-1α signalling early after infection (detected using a *phd3:GFP* transgenic line), which was also observed in infected macrophages at 1dpi (Elks *et al*, 2013), indicating that *il-1β*, alongside Hif-1α signalling, is part of an immediate pro-inflammatory macrophage response. As with Hif-1α, our data indicate that Mm triggered *il-1β* is not sufficient to control infection with subsequent widespread granuloma formation at later stages, however if primed with high *il-1β* and NO via Hif-1α the immune response is boosted leading to lower infection, towards early infection clearance.

We have previously demonstrated that stabilisation of Hif-1α can aid the zebrafish host to control Mm infection, at least in part by priming neutrophils with increased nitrotyrosine generated by the Nos2 enzyme (Elks *et al*, 2013). If the Nos2 enzyme is blocked either pharmacologically or genetically the protective effect of Hif-1α stabilisation is lost (Elks *et al*, 2013). Here, we show that stabilisation of Hif-1α upregulates pro-inflammatory macrophage *il-1β* expression in the absence of an infection challenge. If Il-1β activity is repressed then Hif-1α induced reduction in bacterial burden is abrogated, alongside the Hif-1α dependent increase in NO production. These data show regulation of both Nos2 and Il-1β by Hif-1α, and that Hif-1α driven NO production is partially dependent on Il-1β induction. Both human NOS-2 and IL-1β have HREs (HIF responsive elements) in their promoters and direct regulation by HIF-α signalling has been previously demonstrated *in vitro* (Zhang *et al*, 2006; Charbonneau *et al*, 2007). The link between HIF-1α and IL-1β has been previously demonstrated in murine macrophages via inflammatory activation by succinate, in the absence of infection (Tannahill *et al*, 2013). In a murine model of *Mycobacterium tuberculosis* it was found that HIF-1α is critical for IFN-y–dependent control of M. tuberculosis infection, but it has not previously been demonstrated that HIF-1α is important for innate defense of macrophages against M. tuberculosis (Braverman *et al*, 2016). Our data do not rule out direct regulation of Nos2 by Hif-1α, as blocking Il-1β is likely to have wider spread immune effects, however they do suggest that Nos2 is partially upregulated by Il-1β in the stabilised Hif-1α context. These observations, alongside our finding that blocking Il-1β, primarily observed in macrophages, can block Hif-1α induced neutrophil nitrotyrosine production, indicate a close interplay between macrophages and neutrophils during early mycobacterial infection that is not yet fully understood.

IL-1β is an important pro-inflammatory component and is one of the cytokines that has been shown to be transcriptionally depressed during the 6 hour to 1dpi period of Mm/zebrafish pathogenesis (Benard *et al*, 2016). Although this depression was not detectable using the *il-1β:GFP* line, presumably due to the early transcriptional response post-infection coupled with the stability of the GFP protein, our data indicate that increased *il-1β* transcription due to Hif-1α stabilisation during this early stage of Mm infection is protective to the host. Alongside transcription, the processing of Il-1β by caspases plays a crucial role in immune cell pyroptosis mediated by the inflammasome (Malik & Kanneganti, 2017). Recent findings in the Mm/zebrafish model indicate that neutrophils and macrophages can efficiently phagocytose bacteria and undergo rounds of cell death and re-uptake during the initial days of infection (Hosseini *et al*, 2016). Although here we show a role for early pro-inflammatory *il-1β* transcription during Mm infection, the role of Il-1β processing and inflammasome induced pyroptosis/cell death in these early Mm immune processes remain undetermined.

In conclusion, our data demonstrate an early pro-inflammatory response of Mm infected macrophages *in vivo.* By stabilising Hif-1α, macrophage Il-1β can be primed in the absence of infection and is protective upon Mm infection via neutrophil nitric oxide production. Therapeutic strategies targeting these signalling mechanisms could decrease the level of initial mycobacteria in patients and act to block the development of active TB by reactivation of macrophage pro-inflammatory stimuli. Furthermore, our findings may have important implications in other human infectious diseases in which the pathogen is able to circumvent the proinflammatory immune response to allow its survival and proliferation. Therapies that target host-derived signalling pathways such as these would be beneficial against multidrug resistant strains and could act to shorten the currently long antibiotic therapies required to clear TB from patients.

## Materials and Methods

### Zebrafish and bacterial strains

Zebrafish were raised and maintained on a 14:10-hour light/dark cycle at 28 degrees C, according to standard protocols (Nusslein-Volhard C, 2002), in UK Home Office approved facilities at The Bateson Centre aquaria at the University of Sheffield. Strains used were Nacre (wildtype), *Tg(mpeg1:mCherry-F)ump2Tg, TgBAC(il-1β:eGFP)sh445 Tg(mpeg1:mCherryCAAX)sh378* and *Tg(lyz:Ds-RED2)nz50* (Marjoram *et al*, 2015; Bojarczuk *et al*, 2016; Nguyen-Chi *et al*, 2015; Hall *et al*, 2007).

Mm infection experiments were performed using M. marinum M (ATCC #BAA-535), containing a psMT3-mCherry or psMT3 mCrimson vector (van der Sar *et al*, 2009). Injection inoculum was prepared from an overnight liquid culture in the log-phase of growth resuspended in 2% polyvinylpyrrolidone40 (PVP40) solution (CalBiochem) as previously described (Cui *et al*, 2011). 100-150 colony forming units (CFU) were injected into the caudal vein at 28-30hpf as previously described (Benard *et al*, 2012).

### Generation of *TgBAC(il-1α:GFP)sh445* transgenic and *il-lβ^SH446^/ il-1βř^H446^* mutant zebrafish

An eGFP SV40 polyadenylation cassette inserted at the *il-1β* ATG start site of zebrafish BAC CH-211-147h23 using established protocols (Renshaw *et al*, 2006). Inverted Tol2 elements were inserted into the chloramphenicol coding sequence and the resulting modified BAC was used to generate *TgBAC(il-1 β:eGFP)sh445. il-1-/-(il-10^SH446^/il-10^SH446^)* mutant embryos were generated by CRISPR-Cas9 mediated mutagenesis targeted around an Mwo1 restriction site in the third exon of *il-1β* using the method described by Hruscha et al (2013) and the template sequence 5’-AAAGCACCGACTCGGTGCCACTTTTTCAAGTTGATAACGGACTAGCCTTATTTTA ACTTGCT ATTT CT AGCT CT AAAAC**TGAGCATGTCCAGCACCTC**GGCT AT AGT GA GTCGTATTACGC-3’ (*il-1β* target sequence in bold). PCR with il-1 gF 5’-T AAGG AAAAACT CACTT C-3’ and il-1 gR 5’ATACGTGGACATGCTGAA3’ and subsequent Mwo1 digestion were used for genotyping.

### Morpholino knockdown of *il-1β*

The *il-1bβ*morpholino (Genetools) was used as previously reported (López-Muñoz *et al*, 2011). A standard control morpholino (Genetools) was used as a negative control.

### Confocal microscopy of transgenic larvae

1dpi and 4dpi transgenic zebrafish larvae infected with fluorescent Mm strains were mounted in 0.8-1% low melting point agarose (Sigma-Aldrich) and imaged on a Leica TCS-SPE confocal on an inverted Leica DMi8 base and imaged using 20x or 40x objective lenses.

For quantification purposes acquisition settings and area of imaging (in the caudal vein region) were kept the same across groups. Corrected total cell fluorescence was calculated for each immune-stained cell using Image J as previously described (Elks *et al*, 2013, 2014a).

### Tailfin transection

Inflammation was induced in zebrafish embryos by tail transection at 2 or 3dpf as described previously (Renshaw & Loynes, 2006). Embryos were anesthetised by immersion in 0.168 mg/mL Tricaine (Sigma-Aldrich), and tail transection was performed using a microscalpel (World Precision Instruments).

### qPCR of *il-1β*

SYBR green qPCR was performed on 1dpi Mm infected (or PVP control) embryos as previously described (Van Der Vaart *et al*, 2014). The following primers were used: *il-1β*, accession number NM_212844, forward primer: GAACAGAATGAAGCACATCAAACC, reverse primer: ACGGCACTGAATCCACCAC, *ppial* control, accession number AY391451, forward primer: ACACTGAAACACGGAGGCAAG, reverse primer: CATCCACAACCTTCCCGAACAC.

### Bacterial pixel count

Mm mCherry infected zebrafish larvae were imaged at 4dpi on an inverted Leica DMi8 with a 2.5x objective lens. Brightfield and fluorescent images were captured using a Hammamatsu OrcaV4 camera. Bacterial burden was assessed using dedicated pixel counting software as previously described (Stoop *et al*, 2011).

### RNA injections

Embryos were injected with dominant *hif-1αb* variant RNA at the one cell stage as previously described (Elks *et al*, 2011). *hif-1α* variants used were dominant active (DA) and dominant negative (DN) *hif-1α* (ZFIN: *hif1ab).* Phenol red (PR) (Sigma Aldrich) was used as a vehicle control.

### Anti-nitrotyrosine antibody staining

Larvae were fixed in 4% paraformaldehyde in PBS overnight at 4°C and nitrotyrosine levels were immune-labelled using a rabbit polyclonal anti-nitrotyrosine antibody (Merck Millipore 06-284) and were detected using an Alexa Fluor conjugated secondary antibody (Invitrogen Life Technologies) as previously described (Elks *et al*, 2013, 2014a).

### Statistical analysis

All data were analysed (Prism 7.0, GraphPad Software) using unpaired, two-tailed t-tests for comparisons between two groups and one-way ANOVA (with Bonferonni post-test adjustment) for other data. P values shown are: \**P* < .05, \*\**P* < .01, and \*\*\**P* < .001.

### Funding

This work was supported by a Sir Henry Dale Fellowship jointly funded by the Wellcome Trust and the Royal Society (grant number 105570/Z/14/Z) awarded to (P.M.E.). (S.A.R.) is funded by an MRC Programme Grant (MR/M004864/1). (A.H.M.) is funded by a Smart Mix Program of the Netherlands Ministry of Economic Affairs and the Ministry of Education, Culture and Science. (N.V.O. and H.L.W.) are funded by British Heart Foundation (BHF) project grant (PG/13/80/30443) and Biotechnology and Biological Sciences Research Council (BBSRC) project grant (BB/L000830/1).

## Acknowledgements

The authors would like to thank the Bateson Centre Aquarium Team at the University of Sheffield for fish care. We gratefully thank Georges Lutfalla (Montpellier University) for providing the *Tg(mpeg1 :mCherry-F)ump2Tg line*, Lalita Ramakrishnan (University of Washington, Seattle) for M. marinum strains and Astrid van der Sar (VU University Medical Center, Amsterdam) for the pSMT3-mCherry vector.

## Author Contributions

Conceived and designed the experiments: PME, SAR, NVO. Performed the experiments: PME, NVO, AL. Analysed the data: PME, NVO, HW, AHM, SAR. Wrote the paper: PME, NVO, SAR.

## Conflict of Interest

The authors declare that they have no conflict of interest.

---

**Figure S1. *TgBAC(il-1β:GFP)sh445* recapitulates *il-1β* wholemount *in situ* hybridisation pattern following sterile tailfin transection.**

(A) Wholemount *in situ* hybridisation of *il-1β* in tailfin injured 2dpf embryos. (B) Fluorescent confocal micrographs of *TgBAC(il-1 β:GFP)sh445* expression after tailfin injury. Upper and lower panels show the same individual embryo 0 and 12hpi. (C) Fluorescent confocal micrographs of *TgBAC(il-1 β:GFP)sh445* crossed to *Tg(lyz:Ds-RED2)nz50* labelling neutrophils at 1 hour post wound (1hpw) and 6hpw.

**Figure S2. CRISPR-Cas9 *il-1β SH446* mutant.** (A) Screenshot of Ensembl zebrafish *il-1β* coding sequence with CRISPR-Cas9 target indicated in the fourth exon. (B) DNA alignment of WT *il-1β* sequence and *il-1βSH446* showing the 44 base pair deletion caused by CRISPR-Cas9. (C) Amino acid alignment of WT *il-1β* sequence and *il-1βSH446* with arrowhead showing the premature stop and removal of the putative Il-1β cleavage site. (D) Sequencing trace showing position of CRISPR-Cas9 induced deletion.

**Figure S3. Anti-nitrotyrosine signal is predominantly found in neutrophils after Mm infection.** (A) Example fluorescence confocal z-stacks of the caudal vein region of *Tg(mpx:GFP)i114* embryos (green neutrophils) stained with Alexa-633 labelled anti-nitrotyrosine antibody (cyan), imaged at 1dpi in the presence of Mm mCherry infection (red).

**Supplemental Movie 1. *Il-1β:GFP* expression in activated immune cells after Mm infection.**

(A) Fluorescent confocal videotimelapse of *il-1β:GFP* in immune cells containing Mm infection *(il-1β:GFP* in green and Mm mCrimson in red).

